# Single-cell RNA-seq reveals distinct dynamic behavior of sex chromosomes during early human embryogenesis

**DOI:** 10.1101/382085

**Authors:** Qing Zhou, Taifu Wang, Lizhi Leng, Wei Zheng, Jinrong Huang, Fang Fang, Ling Yang, Jian Wang, Huanming Yang, Fang Chen, Ge Lin, Wen-Jing Wang, Karsten Kristiansen

## Abstract

**Abstract:** *Background:* Several animal and human studies have demonstrated that sex affects kinetics and metabolism during early embryo development. However, the mechanism governing these differences at the molecular level is unknown, warranting a systematic profiling of gene expression in males and females during embryogenesis.

*Findings:* We performed comprehensive analyses of gene expression comparing male and female embryos using available single-cell RNA-sequencing data of 1607 individual cells from 99 human preimplantation embryos, covering development stages from 4-cell to late blastocyst (E2 to E7). Consistent chromosome-wide transcription of autosomes was observed, while sex chromosomes showed significant differences after embryonic genome activation (EGA). Differentially expressed genes (DE genes) in male and female embryos mainly involved in the cell cycle, protein translation and metabolism. The Y chromosome was initially activated by pioneer genes, *RPS4Y1* and *DDX3Y*, while the two X chromosomes in female were widely activated after EGA. Expression of X-linked genes in female significantly declined at the late blastocyst stage, especially in trophectoderm cells, revealing a rapid process of dosage compensation.

*Conclusions:* We observed imbalanced expression from sex chromosomes in male and female embryos during EGA, with dosage compensation occurring first in female trophectoderm cells. Studying the effect of sex differences during human embryogenesis, as well as understanding the mechanism of X chromosome inactivation and its correlation with early miscarriage, will provide a basis for advancing assisted reproductive technology (ART) and thereby improve the treatment of infertility and possibly enhance reproductive health.

## Background

From the moment of fertilization in mammals, the sex of the preimplantation embryo is determined by the spermatozoon carrying either an X or Y chromosome [1, 2]. In recent years, several mammalian and human studies have aimed to molecularly and functionally characterize male and female embryos during *in vitro* development [3, 4]. There are three main aspects of IVF embryos that differ between males and females: 1) patterns of development, including morphology and gene transcription; 2) kinetics and timing of development, including growth rates; 3) mortality during intrauterine development [5]. At the 2-cell stage the percentage of successful culturing differs between male and female mouse embryos [6]. Mouse embryos that carry a Y chromosome develop more quickly *in vitro* than XX embryos [7]. For bovine embryos, addition of the embryonic colony stimulating-factor 2 (CSF2) to the culture medium increases the survival of female embryos at the morula stages, but not male embryos [8]. Moreover, several animal studies have demonstrated that sex affects metabolism during early embryonic development [1, 9, 10]. Thus, much evidence regarding sex differences comes from animal models. For human embryos derived via assisted reproductive technology (ART), it has been reported that embryonic mortality before blastocyst formation is male-biased, as abnormalities occur more frequently in male embryos [11]. In other studies, male IVF embryos are reported to display an increased number of cells and higher metabolic activity than female embryos and develop at a significantly faster rate [12, 13]. While these observations clearly point to developmental differences between male and female embryos, the molecular mechanisms governing these differences prior to the expression of the sex-determine gene *SRY* remain to be established.

At the stage before implantation, sex-specific differences in gene expression become apparent. These have been demonstrated initially in genes that derive from sex chromosomes (at 8-cell stage in mouse [14]), and later in the autosomes (blastocyst in mouse [15]). In bovine embryos, expression of key enzymes involved in establishing genome methylation, as well as histone methylation, is upregulated in male blastocyst compared to their female counterparts [16]. For humans, although Y-chromosome-driven effects have been detected in pluripotent stem cells in a transcriptional study [17], a systematic profiling of gene expression comparing male and female embryos during early development is needed.

Another elusive event in early embryogenesis is X chromosome dosage compensation. In mouse, a period of double X chromosome activation occurs between the 4-cell and 16-cell stages [18, 19]. The burst of transcription from both X chromosomes results in a proteome exhibiting distinct differences between the sexes. Failure to accomplish dosage compensation normally results in early miscarriage and embryonic lethality [20, 21]. In human, it has been reported that X chromosome inactivation occurs in all three lineages of *in vitro* blastocyst embryos on day 7, and that the expression of both X chromosomes is reduced before the random silencing of an entire X chromosome [22]. However, detailed information of the precise temporal activation and inactivation of the X chromosome during early development is still lacking.

The recent development of single-cell sequencing technology has allowed characterizing of individual embryonic cells at multiple levels [23-28] providing comprehensive transcriptional atlases [19, 22, 29, 30]. Here we aimed to determine whether male and female embryos differ in relation to gene expression levels during early development. By analyzing available transcriptome data, we revealed a dynamic pattern of expression for the sex chromosomes and the process of X chromosome dosage compensation in female embryos.

## Data description

To examine whether sex differences already affect transcriptional patterns during early human embryogenesis, we collected available sequencing data on the transcriptome of 1607 individual cells from 99 human preimplantation embryos ranging from the 4-cell stage to late blastocyst [22, 30] (Figure 1A, E2-E7). A total of 3 to 17 embryos and 12 to 466 cells were analyzed per stage (Figure 1B). Single-cell RNA-seq data, processed data and raw reads, on human early embryonic development were downloaded from two publicly available datasets: GSE36552 (78 cells from 4-cell to late blastocyst)[30]; ArrayExpress: E-MTAB-3929 (1529 cells from E3 to E7 embryos)[22]. The DNA methylation data of human embryos was from GSE49828 (covering developmental stages from 4-cell stage to post-implantation) [25].

**Figure1.**
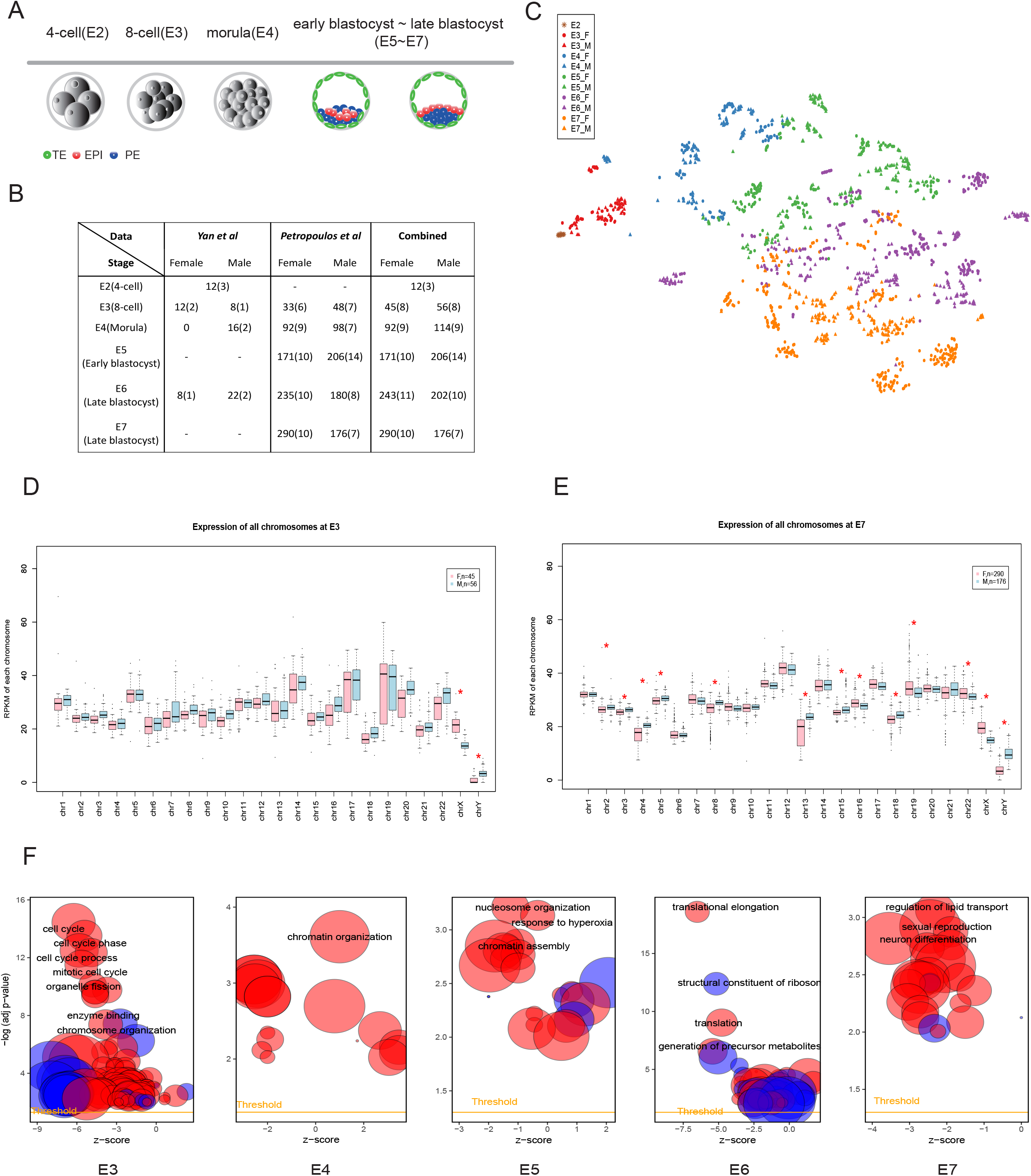
Global transcriptome profiling of male and female embryos reveals differences during development. A) Samples of various development stages included in this study: trophectoderm (TE); epiblast (EPI) and primitive endoderm (PE). B) Table showing the number of male and female cells and embryos used in this study within each embryonic stage. The embryos of 4-cell stage are classified as neither male nor female. C) Two-dimensional t-SNE results of all cells represented by the expression of total genes; Various colors are used to indicate the embryonic day for male (triangle) and female (dot) embryos. D-E) Genome-wide expression per chromosome in the male (light blue) and female (pink) embryos at E3 and E7 stage. Chromosomal RPKM values are calculated as chromosomal reads per kilobase of transcript per million reads mapped (Methods). The significant differences between sexes are defined as P < 10^-5^ (two-sided MWW) and marked with red stars. F) Gene Ontology enrichment results of differentially expressed genes at each stage, representing GO terms for biological processes (red bubble) and molecular function (blue bubble). Summarization of most significant results is listed.x- axis: z-score; y-axis: negative logarithm of the adjusted p-value (provided by DAVID); area of a circle: gene number assigned to the term.

## Analyses

### Transcriptional profiling reveals differences in expression patterns between sex chromosomes during early embryogenesis

We firstly generated the comprehensive transcriptional map of early embryos during these stages. In the result of dimensionality reduction by t-distributed stochastic neighbor embedding (t-SNE), we noticed that the primary segregating factor is the time point of development, not the sex, as samples are clearly classified in agreement with embryonic day (Figure 1C). We then investigated the expression at the chromosome level comparing male and female embryos. Following EGA, differences in gene expression become apparent. At 8-cell stage we observed significant differences in transcription of genes on the sex chromosomes (Figure 1D, p<0.00001, Mann-Whitney-Wilcoxon test), and sex-dependent differences of expression of genes on autosomes become detectable during development, especially when embryos develop to late blastocysts (Figure 1E, Figure S1, p<0.00001, Mann-Whitney-Wilcoxon test).

Next, we performed differential expression analysis comparing female and male cells within embryonic stages. The identified DE genes locate on both sex chromosomes and autosomes for all stages (Table S1, S2). In agreement with previously reported results [22] we found that there is a significant enrichment of DE genes on sex chromosomes (Figure S2, p<0.001, fisher’s exact test). Functional annotation based on Gene Ontology (GO) revealed a stage-specific function for these DE genes (Figure 1G). At the 8-cell stage, these genes are mainly involved in cell cycle control, cell division and chromosomal segregation, and later in the morula stage, DE genes play a role in the chromatin organization. During formation of the blastocyst, sex-dependent differences in gene expression are related to chromatin assembly, translation elongation, and metabolism, and in the late blastocyst stage, DE genes comprise genes involved in regulation of lipid transport and neuron differentiation. All these results indicate that expression differences between males and females are manifest already during early developmental stages of embryogenesis, and thus, regulate various biological processes of development.

### Initial transcription of the Y chromosome is initiated by few pioneer genes

To further investigate the temporal expression patterns of genes on the sex chromosomes, we profiled the expression of Y-linked genes. In total, we detected 27 Y-linked genes exhibiting distinct expression patterns during the early embryonic stages (Figure 2A). For example, transcript from the *PCDH11Y* gene could be detected in E2 embryos before EGA, while down-regulation and transcriptional silencing are observed during later development. Notably, the sex-determining *SRY* gene is as expected not activated at these early stages (Figure S3), reflecting that male gonadal differentiation in the developing embryo first occurs post-implantation [31]. The majority of the Y-linked genes exhibits low levels of expression, but we noticed that two pioneer genes, *RPS4Y1* and *DDX3Y*, exhibit high expression immediately after EGA. These two genes are widely expressed in all male cells, and the sex-specific differences are maintained and even accentuated in the following stages (Figure 2B). Furthermore, we could cluster embryos at the 8-cell stage into two separate groups according to sex solely based on the expression of the *RPS4Y1* gene (Figure 2C). Our analysis reveal that only pioneer genes are highly transcribed during the initial activation of the Y chromosome, and the consistent and high expression in all male cells indicate that the *RPS4Y1* gene could serve as a potential sex-specific marker in human early embryos.

**Figure 2.**
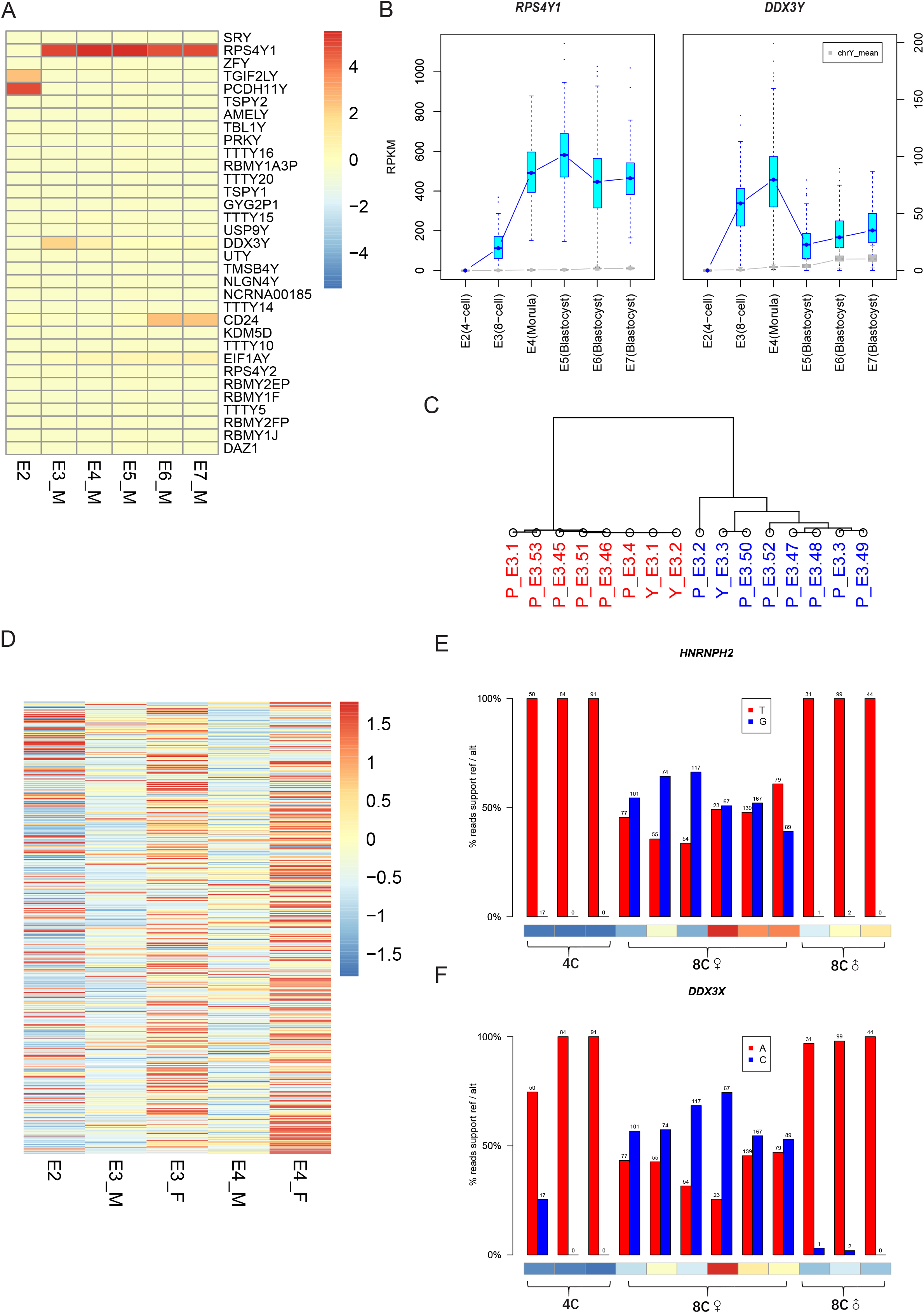
Distinct behavior of sex chromosomes during the process of genome activation. A) Heatmap showing the expression of detected Y-linked genes in the early stages. The mean value within each stage is calculated and scaled to z-score range in [-4,4]; B) Boxplot of the two most highly expressed genes on the Y chromosome indicates high expression in male cells from the 8-cell stage. The boxes with blue color represent the expression of *RPS4X1* and *DDX3Y*, respectively. The mean values of all other Y-linked genes except these two markers are calculated within each cell and drawn as gray boxes. C) Hierarchical clustering for E3 male (blue) and female (red) embryos using the expression of *RPS4Y1* shows a consistent classification pattern. The embryos starting with “Y” are from Yan et al., and the others are from Petropoulos *et al*. D) Heatmap of the average expression of X-linked genes in males and females during the process of genome activation (from E2 to E4). These genes are sorted by their genomic location and the expression is scaled to z-score. E3_M: male embryos at E3; E3_F: female embryos at E3; E4_M: male embryos at E4; E4_F: female embryos at E4. E) Histogram of two representative genes on the X chromosome with biallelic expression in female embryos after EGA. The exact reads number supporting each allele is marked above each bar, and the heatmap under bars present its expression in each cell (with a range from blue to red to show the increase of expression). The two informative loci are *rs41307260* and *rs5963597*.

### Both copies of X chromosome in female are widely activated during EGA

For the X chromosome, we examined the dynamic changes for all expressed genes during the genomic activation process. Still, these X-linked genes, distributed along the entire chromosome, exhibit higher expression in the female than in male embryos after EGA (Figure 2D). Considering the extra copy of the X chromosome in female, we analyzed the allele-specific expression of X-linked genes to determine whether the higher level of expression reflects the activation of both copies. We were able to analyze the allelic expression for each common single nucleotide variant (SNV) present in the dbSNP database within each cell. For example, female embryos at the 8-cell stage show bi-allelic expression of the *HNRNPH2* gene as both a T and a G allele could be identified from the RNA-seq data, whereas the transcript in male embryos harbors only a T allele after EGA (Figure 2E). This is also the case for *DDX3X*, a gene escaping from X-inactivation, with approximate 50% percentage of reads representing expression of the alternative allele in each female cell after EGA (Figure 2F). All the above results demonstrate that all X chromosomes, both in male and female, exhibit wide transcriptional activity during the process of EGA. The transcription of the two copies in females result in an unbalanced dosage between male and female embryos in these early stages.

### Dosage compensation of the X chromosome in female embryos first occurs in the trophectoderm

For adult females, there is a random X chromosome inactivation (XCI) to equalize the expression of X-linked genes with males [32, 33]. As we observed an unbalanced dosage of X chromosome expression comparing males and females in the early embryonic stages, we further focused on the process of dosage compensation in females. To analyze the process of dosage compensation in females, we employed tSNE analysis using only expression of X-linked genes. Despite of the primary classification of development stages, we noticed a sex-specific segregation within each stage (Figure3 A), except for a slight overlap for E7 embryos as they exhibit an overall 70%–85% compensation of X chromosome at that time [22].

**Figure 3.**
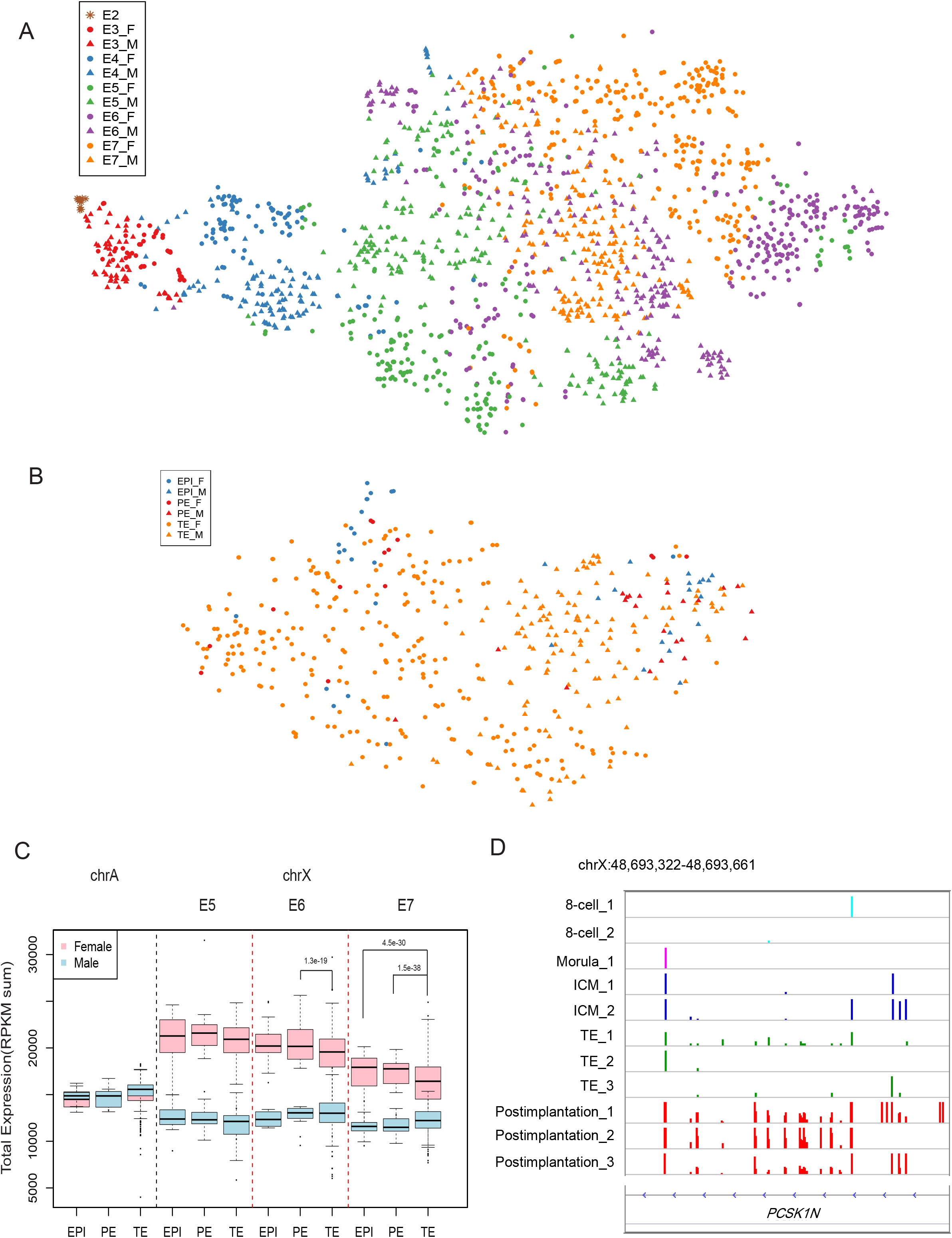
Dosage compensation of the X chromosome firstly occurring in trophectoderm. A) t-SNE results using the expression of X-linked genes during all stages. Various colors indicate the embryonic day for male (triangle) and female (dot) embryos. B) The two -dimension clustering results of E7 embryos, with lineage assignments of EPI (blue), PE (red) and TE (green) for males (triangle) and females (dot). C) Boxplots of X chromosome RPKM sum for individual cells from E5 to E7 blastocysts, with lineage of EPI, PE and TE; an example of chr10 is presented here as negative control of autosomes (chrA); p value, two-sided MWW. D) Integrative genome view (IGV) of the methylation level near to the marker region (chrX:48,693,322-48,693,661) within *PCSK1N* of embryos during development. The height of bars shows the percentage of methylation at each loci, ranging from 0% to 100%.

Since cells are designated as trophectoderm (TE), primitive endoderm (PE), and epiblast (EPI) at the blastocyst stage, we evaluated the X chromosome expression dynamics in the different lineages at this stage. From the tSNE results, we found that overlap between male and female samples could only be detected in TE cells at the E7 stage (Figure 3B). Contrasting the stable expression pattern in male cells (Figure 2A), expression of the X chromosome-linked genes in female tends to be down-regulated with time during the formation of blastocyst, especially in TE cells (Figure 3C, Mann-Whitney-Wilcoxon test). As expected, autosomes show comparable expression in all cells. Expression of X-linked genes declines significantly in TE of E7 embryos, revealing a rapid process of dosage compensation of the X chromosome.

Besides expression profiling, the DNA methylation landscape of specific marker regions on the X chromosome can also reflect the status of gene activation or inactivation [34]. To further support our finding that the first dosage compensation occurs in TE cells, we investigated the DNA methylation level of four reported markers: *AR*, *ZDHHC15*, *SLITRK4* and *PCSK1N* [35]. In total, we collected methylation data for early embryos, including 4-cell, 8-cell, morula, PE (or ICM) and TE from late blastocysts, and post-implantation embryos [25]. As expected, the regions near to *PCSK1N* are hemimethylated in the post-implantation embryos, as one of the X chromosome has completed the inactivation and become methylated at this stage (Figure 3D). Interestingly, we also discovered a low methylation level for these DNA sites in TE cells, comparing with the non-methylated landscape in PE cells (or ICM). Although we could not obtain clear methylation profiles of the other three loci (Figure S4), the specific pattern of *PCSK1N* indicates that methylation as well as inactivation of the X chromosome first occur in female TE cells.

## Discussion

Our study provides to the best of our knowledge the first information on expression differences between male and female IVF embryos during early development. The inclusion of a large number of embryos provides evidence that the transcriptional differences are prominent on sex chromosomes. Our analysis demonstrates distinct transcriptional patterns of genes on the sex chromosomes during early embryogenesis, initial activation of pioneer genes on the Y chromosome and activity of a broad region on the X chromosome. Thus, *RPS4Y1* exhibits high expression at the time of EGA and shows a sex-specific expression pattern. In humans, *RPS4Y1* is one of the variants encoding the ribosomal protein S4 (RPS4), and its paralogous gene *RPS4X* is the first gene on long arm of the X chromosome known to escape from X inactivation [36]. The amino acid differences between the proteins encoded by these two genes result in the generation of two distinct, but functionally equivalent, forms of ribosomes [37]. In contrast to the silencing of the homologous genes in mouse [38], it has been assumed that normal human development requires at least two *RPS4* genes per cell; two *RPS4X* in female cells and one *RPS4X* and one *RPS4Y* in male cells. It has been reported that haploinsufficiency of the ribosomal protein S4 genes may play a role in Turner syndrome [39]. The high transcription of *RPS4Y1* help to balance the dosage between sex [40] at the early stage (Figure S3) as there are two active X chromosomes in female cells after EGA. Thus, expression of *RPS4Y1* may be used as a potential marker to distinguish embryo sex at these stages, earlier than the expression of other sex-determining genes.

The other pioneer gene on the Y chromosome, *DDX3Y*, belongs to the RNA helicase family. The protein encoded by this gene shares high similarity to DDX3X, on the X chromosome, while their functions differ [41]. As a result, activation of this gene in the early stage may lead to a male-specific function, such as neuronal differentiation [42] and translation initiation. In addition, certain part of the ribosome family genes involved in translation elongation and metabolism shows significant differences of expression (Figure S5). As a result, these genes may well regulate protein synthesis and cell signaling pathways. As a sex-ratio bias in relation to of embryonic mortality and growth rates during early development has been reported, all these results may suggest a potential correlation between sex-specific gene expression and the particular behavior of early embryos.

Our study revealed that most X-linked genes become transcriptional active concomitant with completion of EGA in all embryos. In addition, both copies of the X-chromosomes in female are activated. It has been generally assumed that the germline-inactivated X might be passed onto the offspring as in two-cell mouse embryos, where repetitive elements on the paternal X are suppressed [43, 44]. However, *de novo* inactivation of the paternal X chromosome in mouse embryos has been reported [45, 46], with a re-inactivation taking place after the 4-cell stage [19]. For humans, we know that beyond completion of EGA at E4, female cells possess two active X chromosomes [22]. From the comparison of transcription and the allelic expression analysis, our research, for the first time, demonstrates that the two copies of X chromosomes in female are widely activated immediately after genome activation from the 4-cell to the 8-cell stage at E3.

The extensive datasets we investigated in the present study indicate that dosage compensation of the X chromosome first occurred in TE cells. In mice, the imprinted inactivation of the paternal X (Xp) chromosome occurs beyond the 4-cell stage [19]. Inactivity of the Xp is maintained in the TE but is reversed randomly in the ICM of the blastocyst [47, 48]. Key genes, including *Atrx*, which are involved in chromatin remodeling and heterochromatin formation and play a central role in the X-chromosome inactivation process, have been found to be expressed in TE cells, but not in other cell types (EPI). For humans, it has been reported that X chromosome inactivation occurs in all three lineages at E7, and the expression of both X chromosomes is reduced before the random silencing of an entire X chromosome [22]. Our finding of first inactivation in TE cells raises the question as to whether lineage-specific factors, similar to the situation in mouse, can regulate the process of inactivation. Besides, fast inactivation of the X chromosome in TE cells, especially in polar cells (Figure S6) where the first interaction between embryos and uterus occurs during implantation, may result in a balanced dosage between the embryo and the maternal endometrium. Thus, it may be beneficial in relation to implantation as skewed X-chromosome inactivation is associated with recurrent miscarriage [48, 49]. While understanding whether this initial inactivation is paternal imprinted or proceeds randomly is also an important question for the future.

In conclusion, we provide a comprehensive comparison of the transcriptional atlas of male and female human preimplantation embryos and reveal the dynamics of sex chromosomes expression and silencing during embryogenesis for the first time. The precocious X inactivation and decrease in number of TE cells for IVF female embryos may account for the observed preferential female mortality at early stages and the sex ratio in the ART cycle [50]. Studying sex differences during human embryogenesis, as well as understanding the process of X chromosome inactivation and the correlation with early miscarriage, will expand the capabilities of ART and possibly improve the treatment of infertility and enhance reproductive health. In addition, this study of sex differences in early embryos will also provide a basis for further experiments on how environmental impact during early developmental stages can elicit profound and lasting effects that are different in male and female offspring.

## Methods

### Ethical approval

Analyses performed at BGI comprised bioinformatics analysis of public sequencing data, approved by the Institutional Review Board on Bioethics and Biosafety of BGI (IRB 13067).

### Sequencing Data Processing

For RNA-seq data, raw reads were mapped to the human genome (hg19) using TopHat [51, 52] with default settings after removing the low-quality reads. Only uniquely mapped reads were kept for further analysis. Chromosome level expression was counted as chromosomal reads per kilobase of coding region within the chromosome per million mapped reads (RPKM). The gene expression level of raw reads count was calculated by HTSeq [53, 54] and RPKM values were estimated using Cufflinks [51] with the annotation of RefSeq.

### Inference of embryonic sex and cell lineage

Information on sex for each cell and embryo after EGA was classified as previously described [22]. The embryos in the DNA methylation dataset were classified based on the number of detected loci on the Y chromosome, using the average number in oocytes and sperm as a baseline for female and male samples, respectively (Figure S7). The three lineages of cells at the blastocyst stage, as well as the subpopulation group of TE cells, were identified as previously reported [22].

### Sex differential expression analysis

Differential expression analysis was performed for each stage comparing male and female cells. P-values were calculated using DESeq2 [55] and a significant level cut-off of adjusted P<0.05 was used. A cut-off of a 2-fold change in expression was used to define differentially expressed genes. We performed this analysis for each dataset and stage separately, and then combined the results for further annotation. The functional annotation was performed using the Database for Annotation, Visualization and Integrated Discovery (DAVID) [56] Bioinformatics Resource. Gene Ontology terms for each stage were plotted by the GOplot package in R and summarized to a representative term.

### Analyses of allelic expression

The alignment of raw sequencing reads to the human genome was performed by BWA[57], then we employed the function of mpileup in SAMtools [58] to retrieve allelic read counts in the RNA-seq data for common variants in db151[59], and intergenic SNVs were excluded using ANNOVAR [60]. To obtain the total read counts for each site, we run the mpileup program without base quality correction and filtering.

### Statistical analyses

Mann-Whitney-Wilcoxon analyses were performed in R. For identification of DE gene identification, we used an adjusted p value <0.05 as cut-off. In the functional analysis, only GO terms with p<0.01 were included.

## Additional files

**Additional file 1:** Table S1. Differentially analysis results comparing males and females of each embryonic day for two datasets.

**Additional file 2:** Table S2. The distribution of DE genes on each chromosome.

**Additional file 3:** Table S3. Gene Ontology enrichment results of DE genes in each stage.

**Additional file 4:** Figure S1. **(A-C)** Plots of additional stages (E4-E6) for genome-wide expression per chromosome in female (pink box) and male (lightblue box) cells. Chromosomal RPKM values were calculated as chromosomal reads per kilobase of transcript per million reads mapped. The chromosome with a significant difference was marked with a red star if p<10^-5^ in the Mann-Whitney-Wilcoxon test.

**Additional file 5:** Figure S2. **(A-B)** The number of significantly differentially expressed genes on each chromosome comparing males and females at 8-cell stage and late blastocyst in the first dataset (Yan *et al*), stratified by autosomes (green), X chromosome (red) and Y chromosome (blue); **(C-G)** Number of DE genes for each chromosome from E3 to E7 in the other dataset (Petropoulos *et al*). The significant enrichment of DE genes on sex chromosomes was marked with red star (Fisher’s exact test, p<0.001).

**Additional file 6:** Figure S3. **(A)** Boxplots showing expression level of *SRY* from E3 to E7; **(B)** Expression level of *RPS4X* and expression of *RPS4X* and *RPS4Y* in males (light-blue) and females (pink) at each embryonic day(E3-E7). The significant results were marked if p< 0.001 in the Mann-Whitney-Wilcoxon test.

**Additional file 7:** Figure S4. Integrative genome view (IGV) of the DNA methylation patterns of three reported loci (*AR*, *ZDHHC15*, *SLITRK4*) as markers for determining X chromosome inactivation or activation patterns. The height of bars shows the percentage of methylation at each loci, ranging from 0% to 100%. The genomic index: chrX:66,765,297-66,765,584 (*AR*); chrX:74,694,462-74,694,958 (*ZDHHC15*); chrX:142,722,666-142,723,065 (*SLITRK4*).

**Additional file 8:** Figure S5. Heatmap showing the expression of differentially expressed ribosomal genes comparing male and female embryos at E6.

**Additional file 9:** Figure S6. The t-SNE plot of blastocyst cells at E6 **(A)** and E7 **(B)** represented by the expression of X-linked genes. The assignment of subpopulation of cells is indicated as mural (blue), polar (orange) and others (red) for males (triangle) and females (dot).

**Additional file 10:** Figure S7. Bar chart indicating the number of CpG islands detected on the Y chromosome for all embryos in the DNA methylation dataset.

## Author’s contribution

Q.Z., W.J.W and K.K. conceived and designed the study; Q.Z., T.W., J.H., L.Y., F.F., L.L. and W.Z. performed the data analysis; J.W., H.Y., F.C., G.L. oversaw the study; Q.Z., T.W. prepared the figures; Q.Z., T.W., W.J.W. and K.K. wrote and revised the manuscript; all authors reviewed the final version of the manuscript.

## Acknowledgements

The authors thank Chen Ye for data download and management; Huanzi Zhong for constructive advice about the final manuscript.

## Reference

1. Alomar M, Tasiaux H, Remacle S, George F, Paul D, Donnay I. Kinetics of fertilization and development, and sex ratio of bovine embryos produced using the semen of different bulls. Animal Reproduction Science. 2008;107:48–61.

2. Setti AS, Figueira RC, Braga DP, Iaconelli A, Jr., Borges E, Jr. Gender incidence of intracytoplasmic morphologically selected sperm injection-derived embryos: a prospective randomized study. Reprod Biomed Online. 2012;24:420–3.

3. Serdarogullari M, Findikli N, Goktas C, Sahin O, Ulug U, Yagmur E, et al. Comparison of gender-specific human embryo development characteristics by time-lapse technology. Reprod Biomed Online. 2014;29:193–9.

4. Aiken CE, Swoboda PP, Skepper JN, Johnson MH. The direct measurement of embryogenic volume and nucleo-cytoplasmic ratio during mouse pre-implantation development. Reproduction. 2004;128:527–35.

5. Legato MJ. Principles of Gender-Specific Medicine: Gender in the Genomic Era. ELSEVIER:Academic Press. 2017:292–3.

6. Sato E, Xian M, Valdivia RP, Toyoda Y. Sex-linked differences in developmental potential of single blastomeres from in vitro-fertilized 2-cell stage mouse embryos. Horm Res. 1995;44 Suppl 2:4–8.

7. Valdivia RP, Kunieda T, Azuma S, Toyoda Y. PCR sexing and developmental rate differences in preimplantation mouse embryos fertilized and cultured in vitro. Mol Reprod Dev. 1993;35:121–6.

8. Hansen PJ, Dobbs KB, Denicol AC, Siqueira LG. Sex and the preimplantation embryo: implications of sexual dimorphism in the preimplantation period for maternal programming of embryonic development. Cell Tissue Res. 2016;363:237–47.

9. Holm P, Shukri NN, Vajta G, Booth P, Bendixen C, Callesen H. Developmental kinetics of the first cell cycles of bovine in vitro produced embryos in relation to their in vitro viability and sex. Theriogenology. 1998;50:1285–99.

10. Kochhar HP, Peippo J, King WA. Sex related embryo development. Theriogenology. 2001;55:3–14.

11. Orzack SH, Stubblefield JW, Akmaev VR, Colls P, Munne S, Scholl T, et al. The human sex ratio from conception to birth. Proc Natl Acad Sci U S A. 2015;112:E2102–11.

12. Ray PF, Conaghan J, Winston RM, Handyside AH. Increased number of cells and metabolic activity in male human preimplantation embryos following in vitro fertilization. J Reprod Fertil. 1995;104:165–71.

13. Alfarawati S, Fragouli E, Colls P, Stevens J, Gutiérrez-Mateo C, Schoolcraft WB, et al. The relationship between blastocyst morphology, chromosomal abnormality, and embryo gender. Fertility and Sterility. 2011;95:520–4.

14. Lowe R, Gemma C, Rakyan VK, Holland ML. Sexually dimorphic gene expression emerges with embryonic genome activation and is dynamic throughout development. BMC Genomics. 2015;16:295.

15. Kobayashi S, Isotani A, Mise N, Yamamoto M, Fujihara Y, Kaseda K, et al. Comparison of Gene Expression in Male and Female Mouse Blastocysts Revealed Imprinting of the X-Linked Gene at Preimplantation Stages. Current Biology. 2006;16:166–72.

16. Bermejo-Álvarez P, Rizos D, Rath D, Lonergan P, Gutierrez-Adan A. Epigenetic differences between male and female bovine blastocysts produced in vitro. Physiological Genomics. 2008;32:264.

17. Ronen D, Benvenisty N. Sex-dependent gene expression in human pluripotent stem cells. Cell Rep. 2014;8:923–32.

18. Gardner DK, Larman MG, Thouas GA. Sex-related physiology of the preimplantation embryo. Mol Hum Reprod. 2010;16:539–47.

19. Deng Q, Ramskold D, Reinius B, Sandberg R. Single-cell RNA-seq reveals dynamic, random monoallelic gene expression in mammalian cells. Science. 2014;343:193–6.

20. Sullivan AE, Lewis T, Stephenson M, Odem R, Schreiber J, Ober C, et al. Pregnancy outcome in recurrent miscarriage patients with skewed X chromosome inactivation. Obstet Gynecol. 2003;101:1236–42.

21. Lanasa MC, Hogge WA, Kubik C, Blancato J, Hoffman EP. Highly Skewed X-Chromosome Inactivation Is Associated with Idiopathic Recurrent Spontaneous Abortion. The American Journal of Human Genetics. 1999;65:252–4.

22. Petropoulos S, Edsgard D, Reinius B, Deng Q, Panula SP, Codeluppi S, et al. Single-Cell RNA-Seq Reveals Lineage and X Chromosome Dynamics in Human Preimplantation Embryos. Cell. 2016;167:285.

23. Gkountela S, Zhang KX, Shafiq TA, Liao WW, Hargan-Calvopina J, Chen PY, et al. DNA Demethylation Dynamics in the Human Prenatal Germline. Cell. 2015;161:1425–36.

24. Guo F, Yan L, Guo H, Li L, Hu B, Zhao Y, et al. The Transcriptome and DNA Methylome Landscapes of Human Primordial Germ Cells. Cell. 2015;161:1437–52.

25. Guo H, Zhu P, Yan L, Li R, Hu B, Lian Y, et al. The DNA methylation landscape of human early embryos. Nature. 2014;511:606–10.

26. Hou Y, Fan W, Yan L, Li R, Lian Y, Huang J, et al. Genome analyses of single human oocytes. Cell. 2013;155:1492–506.

27. Gao L, Wu K, Liu Z, Yao X, Yuan S, Tao W, et al. Chromatin Accessibility Landscape in Human Early Embryos and Its Association with Evolution. Cell. 2018;173:248–59.e15.

28. Wu J, Xu J, Liu B, Yao G, Wang P, Lin Z, et al. Chromatin analysis in human early development reveals epigenetic transition during ZGA. Nature. 2018;557:256–60.

29. Xue Z, Huang K, Cai C, Cai L, Jiang CY, Feng Y, et al. Genetic programs in human and mouse early embryos revealed by single-cell RNA sequencing. Nature. 2013;500:593–7.

30. Yan L, Yang M, Guo H, Yang L, Wu J, Li R, et al. Single-cell RNA-Seq profiling of human preimplantation embryos and embryonic stem cells. Nat Struct Mol Biol. 2013;20:1131–9.

31. Haqq CM, King CY, Ukiyama E, Falsafi S, Haqq TN, Donahoe PK, et al. Molecular basis of mammalian sexual determination: activation of Mullerian inhibiting substance gene expression by SRY. Science. 1994;266:1494–500.

32. Payer B, Lee JT. X Chromosome Dosage Compensation: How Mammals Keep the Balance. Annual Review of Genetics. 2008;42:733–72.

33. Wutz A. Gene silencing in X-chromosome inactivation: advances in understanding facultative heterochromatin formation. Nat Rev Genet. 2011;12:542–53.

34. Allen RC, Zoghbi HY, Moseley AB, Rosenblatt HM, Belmont JW. Methylation of HpaII and HhaI sites near the polymorphic CAG repeat in the human androgen-receptor gene correlates with X chromosome inactivation. Am J Hum Genet. 1992;51:1229–39.

35. Bertelsen B, Tumer Z, Ravn K. Three new loci for determining x chromosome inactivation patterns. J Mol Diagn. 2011;13:537–40.

36. Fisher EMC, Beer-Romero P, Brown LG, Ridley A, McNeil JA, Lawrence JB, et al. Homologous ribosomal protein genes on the human X and Y chromosomes: Escape from X inactivation and possible implications for turner syndrome. Cell. 1990;63:1205–18.

37. Zinn AR, Alagappan RK, Brown LG, Wool I, Page DC. Structure and function of ribosomal protein S4 genes on the human and mouse sex chromosomes. Molecular and Cellular Biology. 1994;14:2485–92.

38. Zinn AR, Bressler SL, Beer-Romero P, Adler DA, Chapman VM, Page DC, et al. Inactivation of the Rps4 gene on the mouse X chromosome. Genomics. 1991;11:1097–101.

39. Watanabe M, Zinn AR, Page DC, Nishimoto T. Functional equivalence of human X– and Y– encoded isoforms of ribosomal protein S4 consistent with a role in Turner syndrome. Nature Genetics. 1993;4:268.

40. Andrés O, Kellermann T, López-Giráldez F, Rozas J, Domingo-Roura X, Bosch M. RPS4Y gene family evolution in primates. BMC Evolutionary Biology. 2008;8:142-.

41. Rosner A, Paz G, Rinkevich B. Divergent roles of the DEAD-box protein BS-PL10, the urochordate homologue of human DDX3 and DDX3Y proteins, in colony astogeny and ontogeny. Dev Dyn. 2006;235:1508–21.

42. Vakilian H, Mirzaei M, Sharifi Tabar M, Pooyan P, Habibi Rezaee L, Parker L, et al. DDX3Y, a Male-Specific Region of Y Chromosome Gene, May Modulate Neuronal Differentiation. Journal of Proteome Research. 2015;14:3474–83.

43. Cooper DW. Directed genetic change model for X chromosome inactivation in eutherian mammals. Nature. 1971;230:292–4.

44. Huynh KD, Lee JT. Inheritance of a pre-inactivated paternal X chromosome in early mouse embryos. Nature. 2003;426:857–62.

45. Okamoto I, Arnaud D, Le Baccon P, Otte AP, Disteche CM, Avner P, et al. Evidence for de novo imprinted X-chromosome inactivation independent of meiotic inactivation in mice. Nature. 2005;438:369–73.

46. Okamoto I, Otte AP, Allis CD, Reinberg D, Heard E. Epigenetic dynamics of imprinted X inactivation during early mouse development. Science. 2004;303:644–9.

47. Okamoto I, Otte AP, Allis CD, Reinberg D, Heard E. Epigenetic Dynamics of Imprinted X Inactivation During Early Mouse Development. Science. 2004;303:644.

48. Mak W, Nesterova TB, de Napoles M, Appanah R, Yamanaka S, Otte AP, et al. Reactivation of the paternal X chromosome in early mouse embryos. Science. 2004;303:666–9.

49. Dasoula A, Kalantaridou S, Sotiriadis A, Pavlou M, Georgiou I, Paraskevaidis E, et al. Skewed X-Chromosome Inactivation in Greek Women with Idiopathic Recurrent Miscarriage. Fetal Diagnosis and Therapy. 2008;23:198–203.

50. Tarin JJ, Garcia-Perez MA, Hermenegildo C, Cano A. Changes in sex ratio from fertilization to birth in assisted-reproductive-treatment cycles. Reprod Biol Endocrinol. 2014;12:56.

51. Pollier J, Rombauts S, Goossens A. Analysis of RNA-Seq data with TopHat and Cufflinks for genome-wide expression analysis of jasmonate-treated plants and plant cultures. Methods Mol Biol. 2013;1011:305–15.

52. Trapnell C, Pachter L, Salzberg SL. TopHat: discovering splice junctions with RNA-Seq. Bioinformatics. 2009;25:1105–11.

53. Anders S, Pyl PT, Huber W. HTSeq--a Python framework to work with high-throughput sequencing data. Bioinformatics. 2015;31:166–9.

54. Shahriyari L. Effect of normalization methods on the performance of supervised learning algorithms applied to HTSeq-FPKM-UQ data sets: 7SK RNA expression as a predictor of survival in patients with colon adenocarcinoma. Brief Bioinform. 2017.

55. Anders S, Huber W. Differential expression analysis for sequence count data. Genome Biology. 2010;11:R106.

56. Huang DW, Sherman BT, Lempicki RA. Systematic and integrative analysis of large gene lists using DAVID bioinformatics resources. Nature Protocols. 2008;4:44.

57. Li H, Durbin R. Fast and accurate long-read alignment with Burrows–Wheeler transform. Bioinformatics. 2010;26:589–95.

58. Li H. A statistical framework for SNP calling, mutation discovery, association mapping and population genetical parameter estimation from sequencing data. Bioinformatics. 2011;27:2987–93.

59. Sherry ST, Ward MH, Kholodov M, Baker J, Phan L, Smigielski EM, et al. dbSNP: the NCBI database of genetic variation. Nucleic Acids Res. 2001;29:308–11.

60. Wang K, Li M, Hakonarson H. ANNOVAR: functional annotation of genetic variants from high-throughput sequencing data. Nucleic Acids Research. 2010;38:e164–e.

